# Hierarchical Patient-centric Caregiver Network Method for Clinical Outcomes Study

**DOI:** 10.1101/388066

**Authors:** Yoonyoung Park, Panagiotis D. Karampourniotis, Issa Sylla, Amar K. Das

**Affiliations:** IBM Research, Cambridge, MA, USA

## Abstract

In clinical outcome studies, analysis has traditionally been performed using patient-level factors, with minor attention given to provider-level features. However, the nature of care coordination and collaboration between caregivers (providers) may also be important in determining patient outcomes. Using data from patients admitted to intensive care units at a large tertiary care hospital, we modeled the caregivers that provided medical service to a specific patient as patient-centric subnetwork embedded within larger caregiver networks of the institute. The caregiver networks were composed of caregivers who treated either a cohort of patients with particular disease or any patient regardless of disease. Our model can generate patient-specific caregiver network features at multiple levels, and we demonstrate that these multilevel network features, in addition to patient-level features, are significant predictors of length of hospital stay and in-hospital mortality.

## Introduction

Driven by an increased availability of comprehensive healthcare data, interest in predicting patient outcomes and comparing the effectiveness of treatments has been rapidly growing. A majority of health outcomes studies derive analytic models that primarily focus on patient-level factors such as demographic or comorbidity, giving less attention to physicians or other health professionals in charge of patient care. Emphasis on the former assumes that patient characteristics, specific treatments and procedures play more significant roles in determining and explaining clinical outcomes, as compared to healthcare providers, or more broadly, caregiver characteristics. While this may largely be true, for certain medical conditions, coordination and collaboration between multiple caregivers can have a significant role in determining patient outcomes.

In fact, teamwork between providers has been shown to be a strong indicator of quality of care and patient outcomes in hospital settings. Following a four-hour human-based simulator curriculum, Steinemann et al.^1^ observed an immediate improvement in teamwork, speed and completeness of resuscitation among the emergency department (ED) members during the 6 months following the training. The experiments by Morey et al.^2^ employed aviation crew resource management programs within hospital EDs based on the rationale that crew members and caregivers in EDs work in similar environments characterized by time-sensitivity, layered information, and high risk. The experimental groups noticed a 26.5% decrease in clinical errors compared to the control group who did not receive the program. Similarly, Capella et al. demonstrated in their experiment that caregiver teamwork impacts patient outcomes^3^. The experiment formed trauma teams who underwent training sessions and simulations, and resuscitation evaluations. Leadership, communication, situation, and support measurements were improved after the training sessions, as did performance in trauma rooms which ultimately resulted in better patient care.

An increasing number of studies are applying social network analysis to evaluate the importance of caregiver collaboration using healthcare data^4–15^. In many of these studies, a patient-sharing physician network is constructed by assigning an edge (weighted edge) between any two physicians that have treated the same patient (patients). The structures of these emerging patient-sharing networks reveal the impact of institutional boundaries^4^ as well as geographical boundaries^5^. A recent review by Cunningham et al.^15^ concluded that the characteristics of networks are important determinants of quality of care and patient safety. Of particular relevance to our study are the studies by Wang et al.^7^ and by Uddin^8^. In Wang et al.^7^, the authors constructed surgeon-centric collaboration networks based on patients who had undergone knee surgeries using Australian health claims data and examined the association between network topologies and health related outcomes (cost, quality of care). However, the analysis did not take into account patient-level factors that could have affected the study outcomes as well as the patients’ choice of hospital where care was received, raising a possibility for confounding bias. Uddin8 used a similar setting and data to conceptualize a multilevel regression model for evaluating the association between hospital-level variation across 85 hospitals and cost or length of stay for hip replacement surgery. In this study, the author modeled two hospital-level network features, community structure and network density, and used them as clustering variables in the multilevel regression model to show that variation in cluster level affects the coefficients for patient-level features in predicting outcomes.

The importance of caregiver collaboration in patient care demonstrated in prior research suggests that characteristics of caregiver networks can be meaningful predictors of clinical outcomes for patients treated by those in the network. The additive predictive power of such network characteristics beyond that of patients’ demographic or comorbidity features have not been directly evaluated. In our study, we expand the previous efforts by constructing hierarchical patient-sharing caregiver networks in a hospital and using the network features as patient-level predictors of clinical outcomes. We evaluated the importance of network features in predicting the length of hospital stay and in-hospital death.

## Methods

### Data and Study Population

We used Medical Information Mart for Intensive Care III (MIMIC III)^16^, a large public database containing de-identified clinical data from more than 40,000 patients admitted to critical care units at a single tertiary care hospital between 2001 and 2012. MIMIC III expands on MIMIC II, the former version of the dataset which has been widely used in clinical and informatics research. It contains hospitalization-level information such as patient demographics, vital signs, laboratory test results, procedure codes, International Classification of Diseases (ICD) diagnostic codes, medications, intensive care unit (ICU) stays, text notes, de-identified caregiver IDs, and deaths. The dataset is publicly available at https://mimic.physionet.org/.

To examine a group of caregivers treating specific medical conditions, patients admitted for emergency treatment of coronary artery disease or valve disease were identified using Diagnosis-Related Group (DRG) codes. These disease groups were chosen based on the assumption that while regional or institutional variation has been reported^17, 18^, the magnitude of within-institution variation in treatment strategies would be less owing to externally or internally developed guidelines. For the objective of this study, having less variation in patient factors other than the caregiver network was desirable. DRG codes were used instead of ICD codes, because ICD codes associated with each hospitalization are aggregated in summaries without the information about timing of diagnosis. Therefore, a diagnosis of an acute condition can be either the reason for admission, or an event that occurred after the patient was admitted with a different health condition. DRG codes were considered more reliable since they are used for billing purposes and capture the entire episode of hospitalization^19^. Based on DRG codes, we selected 10,378 emergency admissions for treatment of coronary artery or valve disease. Distinct caregiver IDs that provided care during these admissions were identified, which formed the disease-specific caregiver network described below. While all available admission data was used to construct caregiver networks to capture as much information about collaboration as possible, regression analysis was restricted to patients with complete patient-level feature data. In total, 6,621 admission records from 6,368 patients were included in the analysis. Multiple admissions from the same patient were treated as independent admissions, as caregiver assignment is likely independent of who treated the patient in the last admission. The implication of this approach was evaluated in a sensitivity analysis (see Analysis).

### Caregiver Network Construction

In this study framework, patients are treated by a subset of caregivers in a single hospital over their hospitalization period. As each caregiver usually belongs to a single medical specialty department, groups of caregivers who treat a certain patient are more likely to collaborate to treat other patients with similar conditions. Yet, caregivers may also collaborate over patients with other types of conditions. Thus, three levels of collaboration can be identified in this context. The first level is from caregivers treating a specific patient with a particular condition, the second level is an aggregation of collaboration of caregivers treating patients with the same condition, and the third level is an aggregation of all collaborations regardless of patients’ conditions. We used this hierarchical structure to model the collaboration between caregivers, based on the assumption that the extent of collaboration, the resulting network features, and their impact on patient outcomes, would differ depending on which level of network is considered. The caregivers included physicians, nurses, other types (e.g. ‘Pharmacist’, ‘Physician assistants’, etc.), as well as those lacking caregiver description information. Since this information was incomplete in the dataset, those with missing type information were included.

Caregiver networks were constructed by projecting a patient-caregiver network to a caregiver-caregiver network, similar to the method implemented in previous studies^5, 10, 12, 13^. When a group of caregivers treat the same patient in a single admission, edge weights increase by one between all possible pairs of caregivers in that group. Thus, the edge weight between any two nodes in the resulting caregiver network represents the number of unique patient admissions on which the two nodes collaborated. This undirected weighted network captures the collaboration between caregivers across patients. Based on this method, an ‘all-caregiver’ network was constructed using the data from all patients. Similarly, a ‘disease-specific caregiver network’ was constructed based on a subset of data from the cohort of patients with cardiac diseases described above. This level of network captures collaboration for a specific disease group of patients, unlike the all-caregiver network agnostic to patients’ disease groups. Self-loops were not allowed, and caregiver IDs that appeared only once for a single patient were excluded to create a more representative network.

For each patient admission, a ‘subnetwork’ was defined as the subset of nodes (caregivers) in either the all-caregiver or disease-specific caregiver network who treated the patient in that particular admission. Each subnetwork is a fully connected graph. Figure illustrates our approach using simplified graphs. The left panel (A), represents patients (blue nodes) treated by caregivers (green nodes). Each patient (Pt) has a corresponding subnetwork indicated by green ellipses, without showing the edges connecting subnetwork nodes. Patient (Pt) 2, who was treated by only one caregiver, was excluded from further analysis based on patient-sharing caregiver network. The subnetworks are overlaid in the full caregiver network in the right panel (B), weighing each edge by the number of patient-sharing admission episodes. Since caregivers (Cg) 3 and 4 shared two patient admissions, their edge has twice the weight (thicker edge in the Figure) of other edges that shared just one patient.

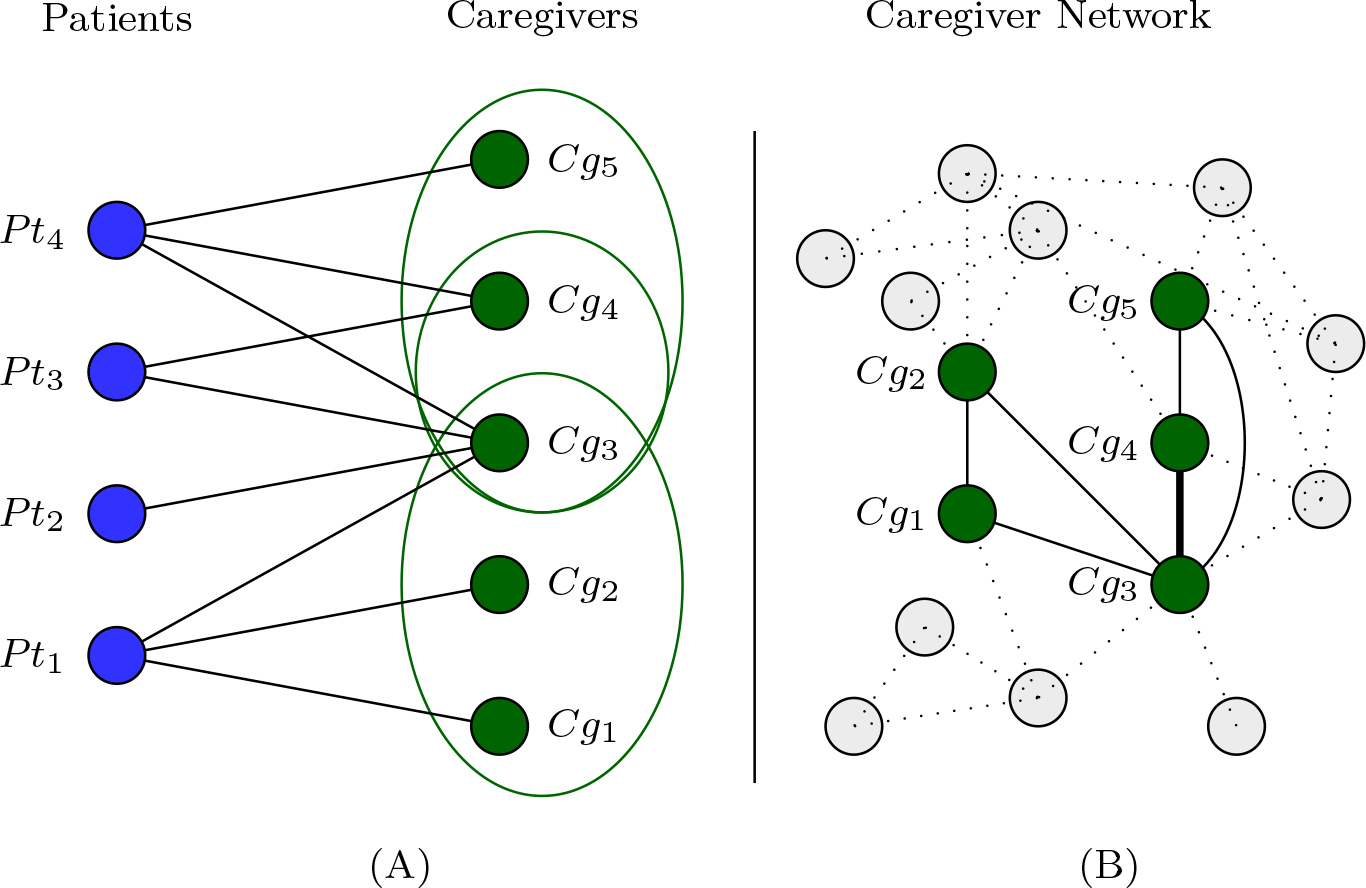
**Networks:** Illustration of patient-centric subnetworks (A) and a caregiver network (B). Patient nodes (Pt) and their caregiver nodes (Cg) generate subnetworks for each patient admission (green ellipses in (A)). Subnetworks are overlaid onto the full caregiver network in panel (B), creating a weighted network of caregivers with edge weight representing the number of shared patient admissions.

### Network Feature Generation

For the constructed network of caregivers *G* := (*V*, *E*), with a set of nodes (vertices) *V* and edges *E*, network level features were generated for each node, and then averaged across the nodes in each patient’s subnetwork to generate patient-level features.

#### Centrality measures

Centrality measures attempt to capture the importance of a node *v* in a network. Degree, betweenness, closeness, and eigenvector centralities are frequently used metrics. Among these, the most basic centrality measure is *degree k*_*i*_ defined by the number of edges that node *i* has in *G*. The set of nodes directly connected to *i* via edges are the neighbors of *i. Betweenness centrality b_i_* is defined by the number of times a node *i* is in the shortest path between any two other nodes in *G*. In hospital settings, it can be considered as the level of control over the flow of information a caregiver has between other caregivers. It is defined as

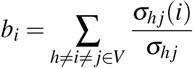

where σ_*hj*_ is the total number of shortest paths from node *h* to *j*, and σ_*hj*_ (*i*) is the number of those paths that pass through node *i*. In our study, we used the edge weights to calculate weighted betweenness centrality using NetworkX. Lastly, *closeness centrality* measures how peripheral a node is, calculated as the average of its distance to all other nodes, while *eigenvector centrality* measures the influence of a node based on its connections to other high influence nodes. In this study, only the degree and betweenness centralities were used to characterize the subnetwork for a given patient as the most conceptually relevant measures, to avoid the collinearity induced by including all measures.

#### Clustering coefficient

Clustering coefficient *c_i_* represents how nodes in a network tend to cluster together, measured by how close the neighboring nodes of *i* are to being a completely connected graph or a clique. In this study, we used the average clustering coefficient (*C*̄)^20^ of each subnetwork as a feature. For the nodes of a subnetwork *S*, the average clustering coefficient is given by

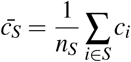

where *n*_*S*_ is the number of nodes in *S* and *c*_*i*_ is the clustering coefficient of node *i* in *G*.

#### Modularity

Modularity *Q* is a metric describing the “strength” of division of a network into clusters, and is often used as the target function in optimization for community detection^21^. It is a comparison of how connected nodes of a particular cluster are to each other than they are to nodes of other clusters and is defined as the difference between the fraction of edges that fall within the clusters and the expected fraction in a random network with equivalent degree distribution and randomly placed edges. High modularity means that the connectivity within clusters is dense compared to connectivity between clusters and indicates a possibility of presence of community. We used a modularity-based metric to determine whether the connectivity of caregivers in a given subnetwork is greater than expected, i.e. whether the subnetwork caregivers tend to work more closely together than what is expected at random.

The weighted, undirected caregiver network is represented by its adjacency matrix *A*. The elements *A_ij_* represent the weight of the edge between *i* and *j*. The modularity metric of a subnetwork *S* is given by

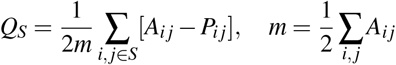

where *A*_*ij*_ and *P*_*ij*_ are the observed and expected at random weights of the edge between nodes *i* and *j*, respectively, and *m* is the sum of all the weights in the network. The expected at random edge weights *P*_*ij*_ are given by the symmetric matrix *P*, constrained by the same total weight *m* as *A*. The strength of a node *r*_*i*_ is defined by the sum of all edge weights of *i*.

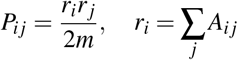

#### Node experience

The level of ‘experience’ of a caregiver node was defined by the number of distinct patient admissions of which the caregiver was a part. While this metric is not directly affected by the neighboring nodes or the network, we hypothesized that the average experience of subnetwork nodes can have an impact on the patient outcomes through more training and professional information gained over time.

### Patient features

After generating network features, patient-level features were obtained from the data. Part of this step utilized MIMIC Code Repository^22^. Demographic and clinical features included age, gender, race/ethnicity, admission location, insurance type, and DRG codes. DRG codes were classified into one of the following categories for ease of analysis: acute coronary artery disease, coronary bypass procedures, cardiac valve replacement procedures, percutaneous coronary interventions (PCI), and other related procedures. Features associated with patient prognosis included use of mechanical ventilation, renal replacement therapy, vasopressor, as well as derived Simplified Acute Physiology Score (SAPS) and noted event of ‘Do Not Resuscitate (DNR)’ or ‘Do Not Intubate (DNI)’ order recorded at the time of ICU admission. If there were more than one ICU stay during a single hospital admission, SAPS values were averaged and other binary features were aggregated to indicate any presence of aforementioned medical events. Elixhauser comorbidity variables, a set of 30 variables used to determine the comorbidity level of inpatients and known to be predictive of hospital length of stay and in-hospital mortality^23^, were used to account for the general health status of a patient. We calculated a single comorbidity score developed by van Walraven et al^24^, using these 30 variables. Patient discharge disposition information was extracted from both structured field and discharge summary notes to identify patients discharged to hospice care.

### Analysis

Descriptive statistics were obtained for patient demographics, clinical characteristics, comorbidity, network degree distribution, and sizes of subnetworks. We focused on two important hospital outcomes: total length of hospital stay (LOS) and in-hospital death. LOS was defined as the time between admission and discharge, and the original LOS value was log-transformed due to the skewed distribution. Continuous variables including network features were standardized by removing the mean and scaling to unit variance. Intermediate features that can change during hospital admission based on the prognosis or health status, such as subnetwork size, were not included in the regression model. In LOS analysis, patients who died in hospital were excluded to prevent potential bias.

Univariate analyses assessed the association between each patient or network feature and either LOS or death. Multivariate regression analyses were conducted in two ways, based on the two levels of caregiver networks as described above (i.e. all-caregiver and disease-specific caregiver networks). First, the features from disease-specific caregiver network were used in the model as they were thought to be the most directly relevant to the outcome in the study cohort. Second, the features from the all-caregiver network were used in the model instead to assess how the collaboration, including care given to other patients outside of the specific disease cohort, captured by all caregiver network, is associated with the outcomes. For LOS, ordinary linear regression models were fitted with 1) patient features only, 2) network features only, and 3) with patient plus network features. For in-hospital death outcome, logistic regression models were fitted using the same three sets of features. The likelihood ratio test was used to compare the fit of different models.

It is possible that terminally ill patients were discharged to hospice before they die, which would make the observed in-hospital mortality an underestimation. As a sensitivity analysis we combined death and discharge to hospice event as an outcome and repeated analysis. Approximately 4% of patients had more than one admission in the data. Possible bias from over-representing those patients was evaluated by randomly selecting one admission from those patients. Lastly, in addition to using a single comorbidity score, we used 30 separate variables and compared the result. When there was a convergence issue with logistic regression models in sensitivity analysis, generalized linear models were used instead with a binomial link function. The presented sensitivity analyses results are based on the disease-specific caregiver network, but the conclusion holds for analyses performed on the all-caregiver network (results not shown).

## Results

From 42,449 patient admissions, we identified 6,621 that met the study criteria including selected cardiac conditions (Table 1). In this cardiac disease-specific cohort, the mean age was 74.1, 34.4% were female, 66.9% were white, and the length of stay was on average 9.1 days among patients who were discharged alive. The average comorbidity score was 3.7 and in-hospital mortality was 6.0% (395 out of 6,621) overall. The DRG code for coronary artery bypass grafting accounted for 36.5% of the patients, followed by percutaneous coronary intervention procedures (29.4%) and valve procedures (18.6%). The most prevalent comorbid disease diagnoses were cardiac arrhythmia (37.4%), diabetes uncomplicated (25.9%), and congestive heart failure (20.3%). Nearly 2.5% of patients received renal replacement therapy, 3.9% had a DNR or DNI order, and more than 60% were mechanically ventilated at some point during hospitalization.

**Table 1.** Selected patient and network characteristics.

| **Patient Characteristics (n = 6,621)** | **Average (Std) or %** |
| --- | --- |
| Age | 74.1 (42.0) |
| Female | 34.4 |
| Race - White | 66.9 |
| Race - African American | 4.1 |
| Race - Asian | 1.5 |
| Race - Hispanic or Latino | 2.4 |
| Race - Others | 25.1 |
| Insurance - Medicare | 57.9 |
| Insurance - Medicaid | 5.2 |
| Insurance - Private | 33.7 |
| Insurance - Others | 3.2 |
| Elixhauser comorbidity score | 3.7 (5.2) |
| SAPS | 18.2 (5.1) |
| Mechanical ventilation | 60.7 |
| Renal replacement therapy | 2.5 |
| Number of vasopressor use | 1.3 |
| DNR or DNI order | 3.9 |
| Length of hospital stay* | 9.1 (6.6) |
| Discharge to hospice* | 0.3 |
| In-hospital death | 6.0 |
| **Network Properties** | **Average (Std)** |
| Degree of nodes in disease-specific network | 354.1 (255.7) |
| Degree of nodes in all caregiver network | 645.3 (453.7) |
| Number of nodes in subnetwork | 14.4 (10.6) |

\*Among patients who were discharged alive; SAPS: Simplified Acute Physiology Score; DNR/DNI: Do Not Resuscitate/Do Not Intubate

The all-caregiver network, constructed irrespective of patient conditions, had 2,310 distinct caregiver nodes with 546,534 patient-sharing edges between the nodes. The disease-specific caregiver network, which is a subset of the all-caregiver network, had 1,303 distinct caregiver nodes with 161,105 edges associated with the patients admitted for heart conditions. The average degree of each caregiver node was 645.3 and 354.1 for the all-caregiver and disease-specific networks, respectively (Table 1). The average size (i.e. number of nodes) of a subnetwork was 14.4, meaning that patients encountered 14 to 15 different caregivers on average while they were hospitalized. The degree distribution of each network is presented in Supplementary Figure.

The results from univariate analyses of patient and network features are presented in Table 2. Overall female gender, admission by referral, private insurance, comorbidity score, SAPS, use of mechanical ventilation, renal replacement therapy, number of vasopressor use, and DRG categories were associated with LOS or risk of death. Among the network features, notably modularity had significantly positive associations with LOS and risk of death, whereas caregiver experience had negative associations with LOS and risk of death. Other network features such as centrality measures and clustering coefficient had differing associations in direction and magnitude with LOS and risk of death.

**Table 2.** Univariate analyses of patient and network features.

| Features | LOS (n = 6226) |  | Death (n = 6621) |  |
| --- | --- | --- | --- | --- |
|  | Coef. | p > \|t\| | Coef. | p > \|t\| |
| Age | 2.83E-04 | 0.11 | 4.39E-03 | < 0.01 |
| Female | 0.07 | < 0.01 | 0.46 | < 0.01 |
| Referral (vs. Urgent<sup>a</sup>) | -0.09 | < 0.01 | -0.29 | 0.02 |
| Transfer (vs. Urgent) | -4.12E-03 | 0.78 | -0.33 | < 0.01 |
| Medicaid (vs. Medicare<sup>b</sup>) | 0.11 | < 0.01 | 0.17 | 0.43 |
| Private (vs. Medicare) | -0.20 | < 0.01 | -0.90 | < 0.01 |
| Self pay (vs. Medicare) | -0.08 | 0.36 | 1.05 | 0.01 |
| Government* (vs. Medicare) | -0.02 | 0.59 | -0.73 | 0.11 |
| Asian (vs. White<sup>c</sup>) | -0.05 | 0.38 | 0.04 | 0.93 |
| African American (vs. White) | 0.07 | 0.07 | 0.39 | 0.08 |
| Hispanic/Latino (vs. White) | 0.04 | 0.39 | -0.93 | 0.07 |
| Others (vs. White) | -0.09 | < 0.01 | 0.26 | 0.02 |
| Acute CAD (vs. CABG<sup>d</sup>) | -0.16 | < 0.01 | 1.57 | < 0.01 |
| PCI (vs. CABG) | -0.64 | < 0.01 | -0.49 | < 0.01 |
| Valve procedures (vs. CABG) | 0.50 | < 0.01 | -0.22 | 0.12 |
| Other procedures (vs. CABG) | 0.19 | < 0.01 | 1.95 | < 0.01 |
| Comorbidity score | 0.16 | < 0.01 | 0.47 | < 0.01 |
| SAPS | 0.27 | < 0.01 | 1.00 | < 0.01 |
| Mechanical ventilation | 0.51 | < 0.01 | 0.42 | < 0.01 |
| Renal replacement therapy | 0.38 | < 0.01 | 1.40 | < 0.01 |
| Number of vasopressor use | 0.14 | < 0.01 | 0.18 | < 0.01 |
| DNR or DNI order | 0.06 | 0.25 | 3.21 | < 0.01 |
| **Disease-specific caregiver network** |  |  |  |  |
| Average degree centrality | -0.10 | < 0.01 | 0.03 | 0.55 |
| Average betweenness centrality | -0.01 | 0.44 | 0.40 | < 0.01 |
| Average clustering coefficient | 0.12 | < 0.01 | -0.01 | 0.83 |
| Modularity | 0.32 | < 0.01 | 0.30 | < 0.01 |
| Average caregiver experience | -0.19 | < 0.01 | -0.37 | < 0.01 |
| **All caregiver network** |  |  |  |  |
| Average degree centrality | -0.13 | < 0.01 | 0.53 | < 0.01 |
| Average betweenness centrality | -0.13 | < 0.01 | 0.07 | 0.12 |
| Average clustering coefficient | 0.15 | < 0.01 | -0.39 | < 0.01 |
| Modularity | 0.29 | < 0.01 | 0.49 | < 0.01 |
| Average caregiver experience | -0.24 | < 0.01 | -0.09 | 0.09 |

<sup>a</sup>Admission type; <sup>b</sup>Insurance type; <sup>c</sup>Race/Ethnicity; <sup>d</sup>DRG category; \*Can be any government insurance; SAPS: Simplified Acute Physiology Score; DNR/DNI: Do Not Resuscitate/Do Not Intubate

Results from regression models including both patient features and network features are presented in Table 3 (showing only network features, see Supplementary Table 1 for all results including patient features). After excluding patients who died in hospital, the linear regression model for LOS with only patient features included had an adjusted *R*^2^ of 0.46, and when disease-specific caregiver network features were included in addition to the patient features, the adjusted *R*^2^ improved to 0.57. All five disease-specific network features were statistically significant predictors of LOS, and the likelihood ratio test comparing the two models suggested that the added network features are meaningful in explaining the length of hospital stay (p < 0.05). With the model predicting LOS built using patient features and all-caregiver network features that captures collaboration for both patients with cardiac diseases and other patients had a similar adjusted *R*^2^ of 0.58, and all network features except for the average betweenness centrality showed significant association with LOS. For predicting in-hospital death in the logistic regression model, none of the disease-specific network features were statistically significant. Among the five all-caregiver network features, degree centrality, clustering coefficient, modularity, and caregiver experience had significant association with in-hospital death. The addition of network features, either from disease-specific or all-caregiver network, improved the prediction model fit based on the likelihood ratio test compared to the model with patient features only.

**Table 3.**
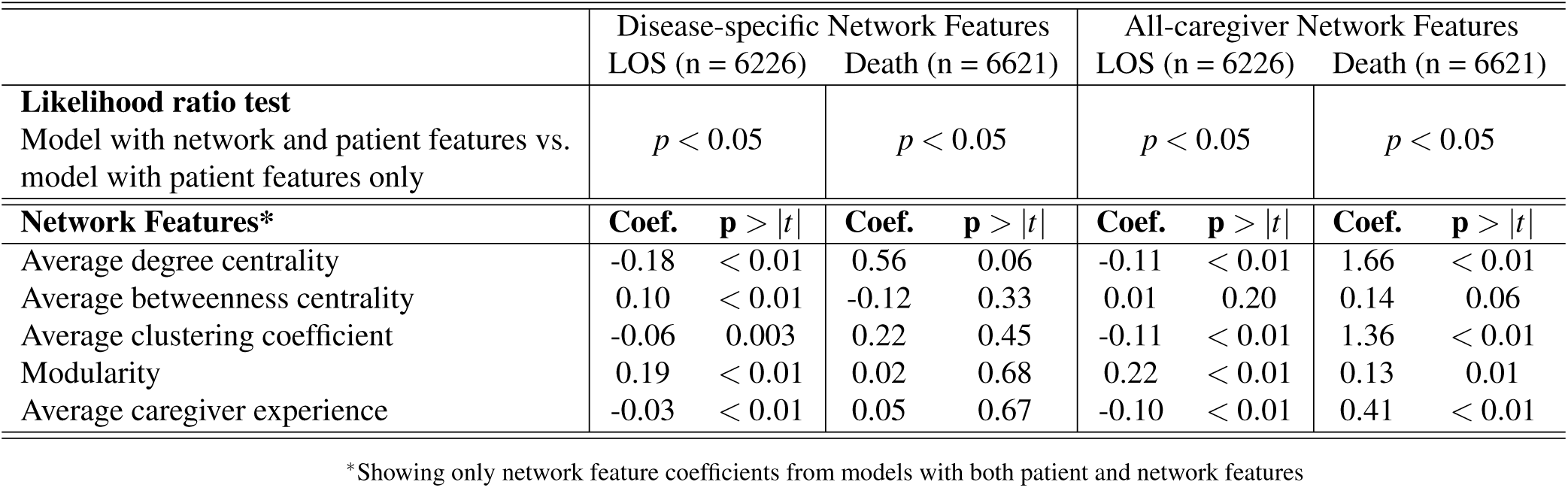
Regression models with both patient and network features.

Sensitivity analysis combining hospice and death as the outcome produced almost identical results, as there were only 18 patients who were identified as discharged to hospice. Similarly, randomly selecting one admission for each patient produced numerically similar results and did not alter the qualitative conclusion. Using 30 disease indicators instead of a combined score to adjust for comorbidity produced numerically different estimates for the regression coefficients but the conclusion from the study remained largely unchanged, that the network features are significant predictors of LOS but not in-hospital mortality (see Supplementary Table 2 for sensitivity analysis results).

## Discussion

In this study, we constructed a hierarchical patient-sharing network to characterize the caregiver collaboration at the patient level and evaluated their association with clinical outcomes. A number of network features were associated with length of stay or in-hospital death outcomes even after adjusting for patient demographic and comorbidity features. Although there was variability in direction and magnitude of the estimated coefficients, the qualitative conclusions were not contradicting between the model using disease-specific caregiver network features and the model using all-caregiver network features, taking statistical precision into account. The study results should be interpreted in a clinical context with a cautionary note since they do not indicate causal effect. For example, in the univariate analysis, average caregiver experience was negatively associated with both LOS and the risk of death. One can imagine a scenario in which the subnetworks comprised of members who have seen greater number of patients are associated with shorter LOS or reduced risk of death through more experience and knowledge. On the other hand, modularity, which depicts the clustering level of the subnetwork compared to a random network, was strongly associated with longer LOS and higher risk of death. The exact clinical interpretation of these network properties needs to be further explored using higher resolution data.

A number of previous studies examined healthcare networks and their characteristics using large datasets. Mandl et al.^9^ modeled both provider-centric and patient-centric constellations using large administrative claims data from the US and provided a detailed characterization of the constellation networks. The study was largely descriptive, did not differentiate between disease types, and did not examine the association between network features and health related outcomes. Landon et al.^10^ built a patient sharing network using Medicare claims data to examine its impact on cost, utilization of service, and quality of care outcomes. The authors improved their previous methods by defining patient-sharing events based on distinct episodes of care, in order to exclude patient sharing for unrelated care. They also used community detection methods based on modularity maximization and adjusted for patient characteristics. But the study aggregated all patient data without focusing on particular diseases or types of care, despite the potential variation in the impact of network features between different diseases. Pollack et al.^6^ created nested networks based on the two specific disease groups: congestive heart failure and diabetes. The authors found that the cost of treatment is lower for patients who are treated by physicians who share more patients. But the network features examined in their study were limited to the density of the networks. Uddin conceptualized ‘patient-centric care network’ in his study8, as a group of physicians who visited the same patient during hospitalization. The approach is different, however, because he used higher level network-level features (community structure and network density, both categorized into five levels) in a multilevel regression model as clustering variables, whereas our study directly adjusted for the network features in the regression models. More importantly, previous studies mostly considered physician collaboration across different hospitals, while our study examined collaboration between different types of caregivers in a single institution. A large part of clinical care is provided by non-physician caregivers, especially for hospitalized patients, therefore supporting our approach as more suitable for the inpatient setting of this study.

We defined a subnetwork from a patient’s perspective, rather than predefining communities in a network and assigning the membership to a patient. In this way, there is a greater flexibility in characterizing each subnetwork in the context of the full network, which is of relevance to inpatient settings where the membership of caregiver groups or ‘communities’ is highly variable. Another strength of the study is our focus on a well-defined clinical scenario that makes interpretation of results straightforward and meaningful. Our approach can be extended to different care settings or different disease cohorts. Of interest for future work is to examine the impact care network features have depending on the patients’ disease(s). Our study is not without limitation, however. Composition of a subnetwork is likely an important factor in characterizing the network, but due to the nature of de-identified data in MIMIC III, we could not take into account the differences between caregiver types. Similarly we could not address the temporality in our analyses, and considering the evolution of networks over time would be one of the future areas of research. As the patients in our study had a number of different DRGs, each with different treatment pathways possible, we did not include in the model the specific treatment that each patient received due to the difficulty of standardizing treatment options for analysis. Instead, we included important procedures with regard to mortality such as mechanical ventilation or renal replacement therapy use, as well as vasopressor use. In addition, adjusting for difference in treatment can potentially influence our ability to observe the effects of network features, since treatment choice can be a downstream effect of network collaboration and act as a mediator of the causal pathway.

Our approach to take caregiver collaboration into account has implications in outcomes research and comparative effectiveness studies, as the quality and nature of care that a patient receives in a hospital may significantly affect patient outcomes. For example, collaboration strength and efficiency may be mediators between treatment effects and clinical outcomes, and particular network compositions may dictate the sequence of procedures a patient receives in a hospital. In addition, our study results suggest that caregiver teamwork and collaboration should be taken into account when evaluating caregiver performance in hospitals. Forming high-functioning teams based on network profiles and characteristics can lead to reduced length of stay and mortality, both of which are important quality measures for hospitals. In conclusion, we show that caregiver network characteristics are important predictors of patient outcome in hospital settings even after adjusting for patient level covariates. The hierarchical network approach is useful in describing the different levels of caregiver collaboration in hospitals, and it can be easily extended to other disease or patient settings.

## Author contributions statement

Y.P. conceived the study, Y.P. and P.K. analyzed the data, and all authors interpreted results and revised the manuscript for intellectual content.

## Additional information

Competing interests: Y.P., P.K., I.S., and A.D. are employees of IBM Research. The study did not receive any specific grant from funding agencies in the public, commercial, or not-for-profit sectors. IBM provided support in the form of salaries for authors Y.P., P.K., I.S., and A.D., but did not have any additional role in the study design, data collection and analysis, decision to publish, or preparation of the manuscript. The specific roles of these authors are articulated in the ‘author contributions’ section. Employment at IBM does not alter our adherence to PLOS ONE policies on sharing data and materials. The authors have no other competing interests to declare.

