## Supplementary for "Hierarchical Patient-centric Caregiver Network Method for Clinical Outcomes Study"

**S1 Figure. Degree distributions of caregiver networks**
**S1 Table 1. Results from multivariate regression models predicting length of stay and in-hospital mortality**
**S1 Table 2. Sensitivity Analysis Results**

**S1 Figure. Degree distributions of caregiver networks**

In a network of caregivers, degree is the number of edges adjacent to each caregiver node.

**a) Distribution of degree of caregiver nodes in all-caregiver network, regardless of patient disease type. Average degree was 645.3 with a standard deviation of 453.7.**

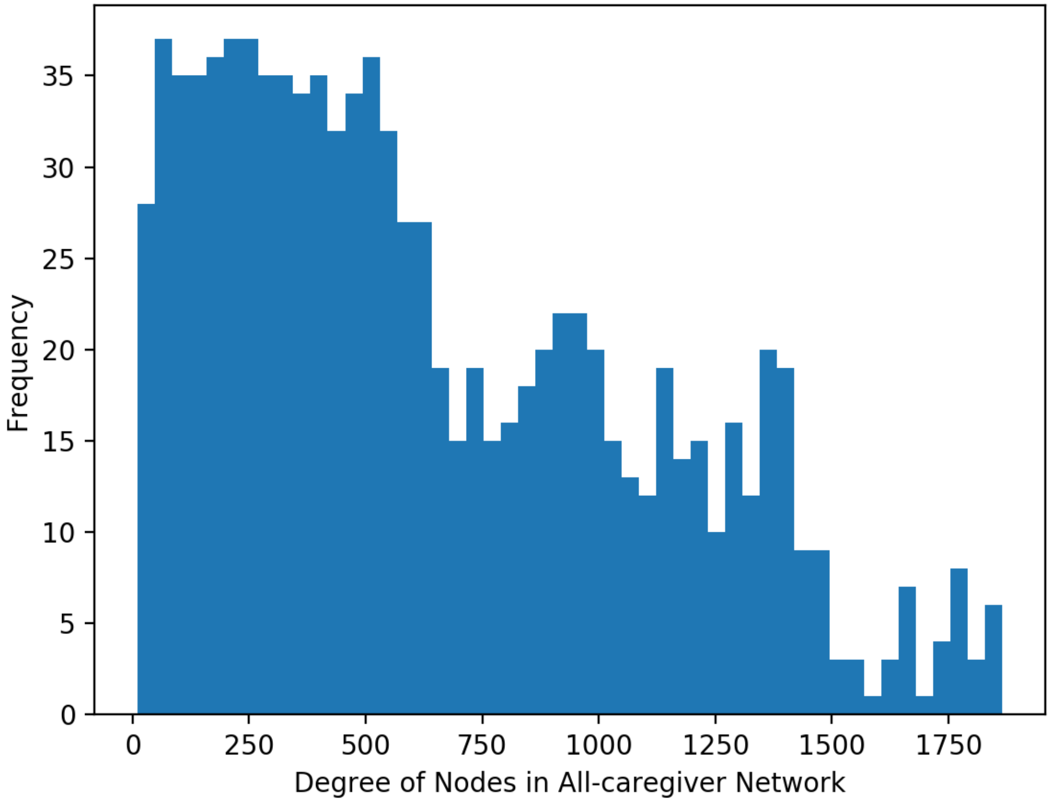

**b) Distribution of degree of caregiver nodes in disease-specific caregiver network for patients with coronary heart disease or valve disease. Average degree was 354.1 with a standard deviation of 255.7.**

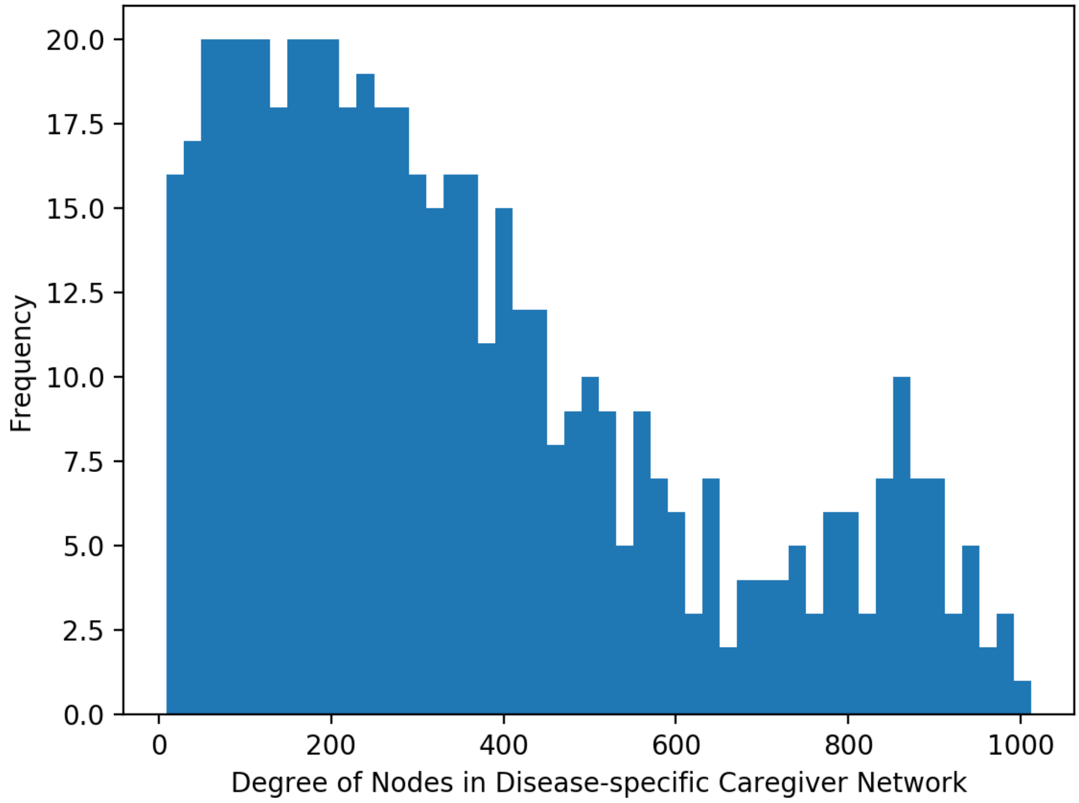

**c) Distribution of subnetwork size, i.e. the number of different caregivers that provided care to a patient during a hospitalization period. The average size of a subnetwork was 14.4, meaning that patients encountered 14 to 15 different caregivers on average while they were hospitalized.**

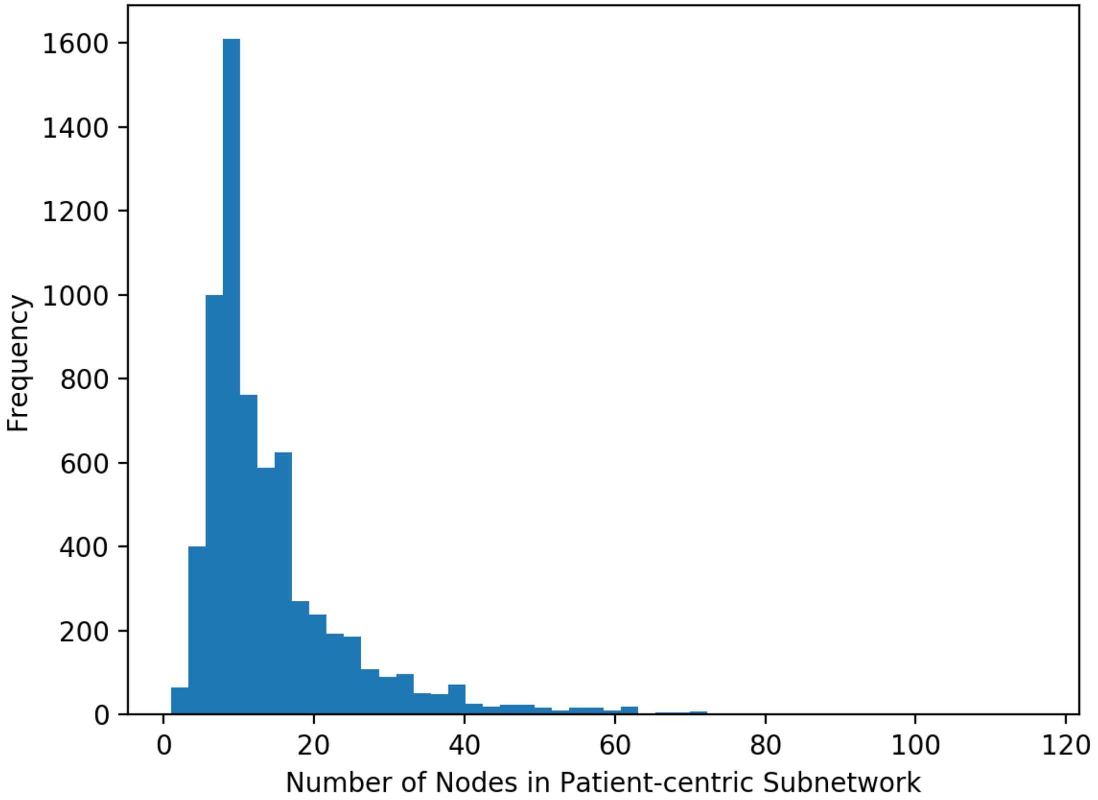

**S1 Table 1. Results from multivariate regression models predicting length of stay and in-hospital mortality**

| | Disease-specific | | | | All-caregiver | | | |
|---|---|---|---|---|---|---|---|---|
| | LOS | | Death | | LOS | | Death | |
| | Coef. | p >\|t\| | Coef. | p >\|t\| | Coef. | p >\|t\| | Coef. | p >\|t\| |
| Age | 3E-04 | 0.02 | -3E-04 | 0.79 | 4E-04 | 0.01 | 1E-04 | 0.91 |
| Female | 0.01 | 0.48 | -0.07 | 0.62 | 0.01 | 0.26 | -0.07 | 0.62 |
| Admission[a] - Referral | -0.14 | < 0.01 | -0.22 | 0.23 | -0.15 | < 0.01 | -0.15 | 0.42 |
| Admission - Transfer | -0.06 | < 0.01 | -0.30 | 0.06 | -0.07 | < 0.01 | -0.26 | 0.10 |
| Insurance[b]- Government | 0.07 | 0.03 | < 0.01 | 1.00 | 0.07 | 0.02 | 0.07 | 0.89 |
| Insurance - Medicaid | 0.08 | < 0.01 | 0.41 | 0.14 | 0.08 | < 0.01 | 0.34 | 0.24 |
| Insurance - Private | -0.07 | < 0.01 | -0.05 | 0.76 | -0.07 | < 0.01 | -0.05 | 0.77 |
| Insurance - Self Pay | 0.04 | 0.46 | 1.43 | 0.01 | 0.04 | 0.52 | 1.23 | 0.02 |
| Race/Ethnicity[c] - Asian | -0.06 | 0.14 | -0.19 | 0.72 | -0.06 | 0.16 | -0.21 | 0.71 |
| Race/Ethnicity - African American | 0.02 | 0.50 | -0.23 | 0.45 | 0.02 | 0.54 | -0.18 | 0.55 |
| Race/Ethnicity - Hispanic/Latino | -0.05 | 0.13 | -0.95 | 0.10 | -0.03 | 0.30 | -0.79 | 0.17 |
| Race/Ethnicity - Others | -0.05 | < 0.01 | 0.49 | < 0.01 | -0.05 | < 0.01 | 0.54 | < 0.01 |
| DRG[d] - Acute CAD | -0.30 | < 0.01 | 3.40 | < 0.01 | -0.31 | < 0.01 | 2.74 | < 0.01 |
| DRG - Other procedures | -0.10 | < 0.01 | 3.04 | < 0.01 | -0.13 | < 0.01 | 2.54 | < 0.01 |
| DRG - PCI | -0.45 | < 0.01 | 1.95 | < 0.01 | -0.47 | < 0.01 | 1.39 | < 0.01 |
| DRG - Valve procedures | 0.12 | < 0.01 | 1.34 | < 0.01 | 0.14 | < 0.01 | 1.32 | < 0.01 |
| Comorbidity Score | 0.09 | < 0.01 | 0.19 | < 0.01 | 0.10 | < 0.01 | 0.19 | < 0.01 |
| SAPS | 0.05 | < 0.01 | 0.94 | < 0.01 | 0.05 | < 0.01 | 0.99 | < 0.01 |
| Mechanical Ventilation | 0.08 | < 0.01 | -0.21 | 0.29 | 0.04 | 0.01 | -0.25 | 0.20 |
| Renal Replacement Therapy | 0.09 | 0.01 | 0.19 | 0.50 | 0.09 | 0.01 | 0.24 | 0.40 |
| DNR or DNI order | 0.09 | 0.01 | 2.20 | < 0.01 | 0.08 | 0.02 | 2.16 | < 0.01 |
| Number of Vasopressor Use | 0.01 | < 0.01 | 0.20 | < 0.01 | 0.01 | < 0.01 | 0.18 | < 0.01 |
| Average betweenness centrality | 0.10 | < 0.01 | -0.12 | 0.33 | 0.01 | 0.20 | 0.14 | 0.06 |
| Average degree centrality | -0.18 | < 0.01 | 0.56 | 0.06 | -0.11 | < 0.01 | 1.66 | < 0.01 |
| Average clustering coefficients | -0.06 | < 0.01 | 0.22 | 0.45 | -0.11 | < 0.01 | 1.36 | < 0.01 |
| Average node experience | -0.03 | < 0.01 | 0.05 | 0.67 | -0.10 | < 0.01 | 0.41 | < 0.01 |
| Modularity | 0.19 | < 0.01 | 0.02 | 0.68 | 0.22 | < 0.01 | 0.13 | 0.01 |

[a] Compared to Emergency Admission; [b] Compared to Medicare; [c] Compared to White; [d] Compared to CABG

LOS: length of stay; CAD: coronary artery disease; PCI: percutaneous coronary intervention; CABG: coronary artery bypass graft; SAPS: Simplified Acute Physiology Score; DNR/DNI: Do Not Resuscitate/Do Not Intubate

**S1 Table 2. Sensitivity Analysis Results**

Sensitivity analyses were conducted based on disease-specific caregiver network to examine the robustness of study results. Results presented below are from multivariable regression models predicting either length of stay or in-hospital death.

**a) Combining discharge to hospice and in-hospital death as an alternative mortality outcome variable**

| | LOS | | Death | |
|---|---|---|---|---|
| | Coef. | p >\|t\| | Coef. | p >\|t\| |
| Age | 3E-04 | 0.02 | -7E-04 | 0.60 |
| Female | 0.01 | 0.53 | -0.03 | 0.80 |
| Admission[a] - Referral | -0.14 | < 0.01 | -0.16 | 0.36 |
| Admission - Transfer | -0.06 | < 0.01 | -0.25 | 0.11 |
| Insurance[b] - Government | 0.07 | 0.03 | 0.35 | 0.44 |
| Insurance - Medicaid | 0.08 | < 0.01 | 0.39 | 0.15 |
| Insurance - Private | -0.07 | < 0.01 | -0.05 | 0.75 |
| Insurance - Self Pay | 0.04 | 0.46 | 1.41 | 0.01 |
| Race/Ethnicity[c] - Asian | -0.05 | 0.18 | -0.04 | 0.94 |
| Race/Ethnicity - African American | 0.02 | 0.51 | -0.26 | 0.38 |
| Race/Ethnicity - Hispanic/Latino | -0.05 | 0.13 | -0.76 | 0.14 |
| Race/Ethnicity - Others | -0.05 | < 0.01 | 0.49 | < 0.01 |
| DRG[d] - Acute CAD | -0.30 | < 0.01 | 3.35 | < 0.01 |
| DRG - Other procedures | -0.10 | < 0.01 | 2.96 | < 0.01 |
| DRG - PCI | -0.45 | < 0.01 | 1.80 | < 0.01 |
| DRG - Valve procedures | 0.12 | < 0.01 | 1.29 | < 0.01 |
| Comorbidity Score | 0.09 | < 0.01 | 0.21 | < 0.01 |
| SAPS | 0.05 | < 0.01 | 0.91 | < 0.01 |
| Mechanical Ventilation | 0.08 | < 0.01 | -0.30 | 0.12 |
| Renal Replacement Therapy | 0.08 | 0.02 | 0.20 | 0.48 |
| DNR or DNI order | 0.11 | < 0.01 | 2.34 | < 0.01 |
| Number of Vasopressor Use | 0.01 | < 0.01 | 0.18 | < 0.01 |
| Average betweenness centrality | 0.10 | < 0.01 | -0.10 | 0.42 |
| Average degree centrality | -0.19 | < 0.01 | 0.40 | 0.16 |
| Average clustering coefficients | -0.06 | < 0.01 | 0.08 | 0.77 |
| Average node experience | -0.03 | < 0.01 | 0.02 | 0.87 |
| Modularity | 0.19 | < 0.01 | 0.04 | 0.54 |

[a] Compared to Emergency Admission; [b] Compared to Medicare; [c] Compared to White; [d] Compared to CABG

LOS: length of stay; CAD: coronary artery disease; PCI: percutaneous coronary intervention; CABG: coronary artery bypass graft; SAPS: Simplified Acute Physiology Score; DNR/DNI: Do Not Resuscitate/Do Not Intubate

**b) Randomly selecting one admission from each patient (n=6,368)**

| | LOS | | Death | |
|---|---|---|---|---|
| | Coef. | p >\|t\| | Coef. | p >\|t\| |
| Age | 3E-04 | 0.02 | -6E-04 | 0.66 |
| Female | 0.01 | 0.55 | -0.08 | 0.54 |
| Admission[a] - Referral | -0.14 | < 0.01 | -0.21 | 0.25 |
| Admission - Transfer | -0.06 | < 0.01 | -0.28 | 0.08 |
| Insurance[b] - Government | 0.05 | 0.14 | 0.04 | 0.94 |
| Insurance - Medicaid | 0.07 | < 0.01 | 0.44 | 0.11 |
| Insurance - Private | -0.08 | < 0.01 | -0.11 | 0.53 |
| Insurance - Self Pay | 0.03 | 0.62 | 1.42 | 0.01 |
| Race/Ethnicity[c] - Asian | -0.06 | 0.16 | -0.39 | 0.50 |
| Race/Ethnicity - African American | 2.2E-03 | 0.93 | -0.29 | 0.35 |
| Race/Ethnicity - Hispanic/Latino | -0.03 | 0.43 | -0.77 | 0.17 |
| Race/Ethnicity - Others | -0.05 | < 0.01 | 0.46 | < 0.01 |
| DRG[d] - Acute CAD | -0.30 | < 0.01 | 3.41 | < 0.01 |
| DRG - Other procedures | -0.11 | < 0.01 | 3.05 | < 0.01 |
| DRG - PCI | -0.46 | < 0.01 | 1.92 | < 0.01 |
| DRG - Valve procedures | 0.12 | < 0.01 | 1.31 | < 0.01 |
| Comorbidity Score | 0.09 | < 0.01 | 0.19 | < 0.01 |
| SAPS | 0.06 | < 0.01 | 0.96 | < 0.01 |
| Mechanical Ventilation | 0.07 | < 0.01 | -0.27 | 0.17 |
| Renal Replacement Therapy | 0.10 | 0.01 | 0.32 | 0.28 |
| DNR or DNI order | 0.10 | 0.01 | 2.22 | < 0.01 |
| Number of Vasopressor Use | 0.01 | < 0.01 | 0.19 | < 0.01 |
| Average betweenness centrality | 0.10 | < 0.01 | -0.15 | 0.24 |
| Average degree centrality | -0.18 | < 0.01 | 0.63 | 0.03 |
| Average clustering coefficients | -0.06 | < 0.01 | 0.29 | 0.32 |
| Average node experience | -0.03 | < 0.01 | 0.05 | 0.69 |
| Modularity | 0.19 | < 0.01 | 0.05 | 0.43 |

[a] Compared to Emergency Admission; [b] Compared to Medicare; [c] Compared to White; [d] Compared to CABG

LOS: length of stay; CAD: coronary artery disease; PCI: percutaneous coronary intervention; CABG: coronary artery bypass graft; SAPS: Simplified Acute Physiology Score; DNR/DNI: Do Not Resuscitate/Do Not Intubate

**c) Using 29* Elixhauser comorbidity variables instead of single comorbidity score value variable**

| | LOS | | Death | |
|---|---|---|---|---|
| | Coef. | p >\|t\| | Coef. | p >\|t\| |
| Age | 3E-04 | 0.05 | 3E-04 | 0.83 |
| Female | 0.01 | 0.37 | -0.06 | 0.66 |
| Admission[a] - Referral | -0.14 | < 0.01 | -0.22 | 0.23 |
| Admission - Transfer | -0.06 | < 0.01 | -0.28 | 0.09 |
| Insurance[b] - Government | 0.06 | 0.04 | -0.35 | 0.56 |
| Insurance - Medicaid | 0.09 | < 0.01 | 0.04 | 0.89 |
| Insurance - Private | -0.05 | < 0.01 | -0.22 | 0.22 |
| Insurance - Self Pay | 0.03 | 0.57 | 1.05 | 0.07 |
| Race/Ethnicity[c] - Asian | -0.04 | 0.26 | -0.01 | 0.98 |
| Race/Ethnicity - African American | 0.01 | 0.66 | -0.22 | 0.48 |
| Race/Ethnicity - Hispanic/Latino | -0.04 | 0.17 | -0.88 | 0.13 |
| Race/Ethnicity - Others | -0.04 | < 0.01 | 0.42 | 0.01 |
| DRG[d] - Acute CAD | -0.32 | < 0.01 | 3.50 | < 0.01 |
| DRG - Other procedures | -0.11 | < 0.01 | 3.05 | < 0.01 |
| DRG - PCI | -0.45 | < 0.01 | 2.05 | < 0.01 |
| DRG - Valve procedures | 0.08 | < 0.01 | 1.79 | < 0.01 |
| SAPS | 0.04 | < 0.01 | 0.96 | < 0.01 |
| Mechanical Ventilation | 0.08 | < 0.01 | -0.34 | 0.10 |
| Renal Replacement Therapy | 0.03 | 0.47 | 0.45 | 0.17 |
| DNR or DNI order | 0.08 | 0.02 | 2.35 | < 0.01 |
| Number of Vasopressor Use | 0.01 | < 0.01 | 0.20 | < 0.01 |
| Average betweenness centrality | 0.11 | < 0.01 | -0.14 | 0.29 |
| Average degree centrality | -0.18 | < 0.01 | 0.62 | 0.04 |
| Average clustering coefficients | -0.05 | 0.01 | 0.25 | 0.39 |
| Average node experience | -0.02 | 0.02 | 0.09 | 0.43 |
| Modularity | 0.18 | < 0.01 | -1.8E-03 | 0.98 |
| Congestive heart failure | 0.11 | < 0.01 | -0.14 | 0.42 |
| Cardiac arrhythmias | 0.09 | < 0.01 | 0.01 | 0.93 |
| Valvular disease | 0.04 | 0.02 | -0.80 | < 0.01 |
| Pulmonary circulation | 0.08 | < 0.01 | -0.06 | 0.80 |
| Peripheral vascular | 0.06 | < 0.01 | 0.07 | 0.66 |
| Hypertension | 0.04 | 0.09 | -0.28 | 0.30 |
| Paralysis | 0.17 | < 0.01 | -0.35 | 0.50 |
| Other neurological disease | 0.12 | < 0.01 | 0.92 | < 0.01 |
| Chronic pulmonary | 0.04 | < 0.01 | -0.02 | 0.89 |

| | | | | |
|---|---|---|---|---|
| Diabetes uncomplicated | 0.01 | 0.26 | 0.09 | 0.55 |
| Diabetes complicated | 0.11 | < 0.01 | 0.01 | 0.96 |
| Hypothyroidism | -0.06 | 0.35 | -0.38 | 0.72 |
| Renal failure | 0.06 | 0.08 | 0.14 | 0.72 |
| Liver disease | 0.05 | 0.32 | 1.20 | 0.02 |
| Peptic ulcer disease | -0.09 | 0.68 | -19.80 | 1.00 |
| AIDS | -3.1E-16 | < 0.01 | 1.4E-07 | 1.00 |
| Lymphoma | 0.06 | 0.35 | -21.37 | 1.00 |
| Metastatic cancer | 0.22 | < 0.01 | 0.63 | 0.16 |
| Solid tumor | 0.09 | 0.02 | -0.17 | 0.76 |
| Rheumatoid arthritis | 0.06 | 0.54 | 0.33 | 0.73 |
| Coagulopathy | 0.06 | < 0.01 | 0.83 | < 0.01 |
| Obesity | -4E-03 | 0.85 | 0.85 | < 0.01 |
| Weight loss | 0.41 | < 0.01 | -0.49 | 0.39 |
| Fluid electrolyte | 0.07 | < 0.01 | 0.74 | < 0.01 |
| Blood loss anemia | 1.0E-18 | 0.93 | 1.7E-08 | 1.00 |
| Deficiency anemias | 0.07 | < 0.01 | -0.98 | < 0.01 |
| Alcohol abuse | -0.01 | 0.71 | 0.44 | 0.23 |
| Drug abuse | 0.18 | < 0.01 | -0.66 | 0.34 |
| Psychoses | 0.03 | 0.40 | -0.18 | 0.73 |

*There are 30 Elixhauser variables, but depression was not included because there was no patient with depression in the study data.

[a] Compared to Emergency Admission; [b] Compared to Medicare; [c] Compared to White; [d] Compared to CABG

LOS: length of stay; CAD: coronary artery disease; PCI: percutaneous coronary intervention; CABG: coronary artery bypass graft; SAPS: Simplified Acute Physiology Score; DNR/DNI: Do Not Resuscitate/Do Not Intubate; AIDS: acquired immunodeficiency syndrome
